# Reassembling a cannon in the DNA defense arsenal: genetics of StySA, a BREX phage exclusion system in *Salmonella* lab strains

**DOI:** 10.1101/2021.11.15.468586

**Authors:** Julie Zaworski, Oyut Dagva, Julius Brandt, Chloé Baum, Laurence Ettwiller, Alexey Fomenkov, Elisabeth A. Raleigh

## Abstract

Understanding mechanisms that shape horizontal exchange in prokaryotes is a key problem in biology. A major limit on DNA entry is imposed by restriction-modification (RM) processes that depend on the pattern of DNA modification at host-specified sites. In classical RM, endonucleolytic DNA cleavage follows detection of unprotected sites on entering DNA. Recent investigation has uncovered BREX systems, RM-like activities that employ host protection by DNA modification but replication arrest without evident nuclease action on unmodified phage DNA. We show that the historical *stySA* RM locus of *Salmonella enterica* sv Typhimurium is a BREX homolog. The *stySA29* allele of the hybrid strain LB5000 carries a mutated version of the ancestral LT2 BREX system. Surprisingly, both a restriction and a methylation defect are observed for this lineage despite lack of mutations in *brxX*, the modification gene homolog. Instead, flanking genes *pglZ* and *brxC* each carry multiple mutations (μ) in C-terminal domains. To avoid plasmid artifacts and potential stoichiometric interference, we chose to investigate this system *in situ*, replacing the mutated *pglZμ* and *brxCμ* genes with wild type (WT). PglZ-WT supports methylation in the presence of either BrxCμ or BrxC-WT but not in the presence of a deletion/insertion allele, Δ*brxC::cat*. Restriction of phage L requires both BrxC-WT and PglZ-WT, implicating the BrxC C-terminus specifically in restriction activity. Disruption of four other CDS with *cat* cassettes still permitted modification, suggesting that BrxC, PglZ and BrxX are principal components of the modification activity. BrxL is required for restriction only. A partial disruption of *brxL* disrupts transcription globally.

## Introduction

Transfer of genes from one organism to another shapes ecological capacities in microbiomes on both short and long-term timescales. Thus, mechanisms that limit or promote such transfer are of fundamental interest. Ecologic interactions with phage play a major role in host colonization by prokaryotes [1, 2]. Prokaryote defenses against phages are of particular interest [3, 4], particularly as therapeutic uses are contemplated [5]. Microbiome and metagenome studies have led to a renaissance in the study of phage-host interaction [6, 7].

Host defensive processes include restriction-modification (RM) systems as major contributors[8]. These distinguish the host DNA from foreign invaders using the pattern of DNA modification; DNA with the wrong modification pattern is rejected [9, 10]. DNA modification by bacterial hosts is typically carried out by methylation of adenine or cytosine bases. Recently, sequence-specific protective phosphorothioation of the DNA backbone has emerged as an important biological phenomenon [11]. In most familiar cases of RM systems, protection is conferred by methylation of a particular base within a specific sequence motif, while rejection consists of endonucleolytic cleavage of both strands in response to the presence of unmethylated specific sites. Such cleavage leads to very rapid interruption of the phage development program.

Both defensive and epigenetic processes can involve DNA modification states, so taxa with no DNA modification are extremely rare. Epigenetic regulation is important in the life of the cell, and often the relevant genes are fixed in a lineage, while defense functions are diverse [12, 13].

A further result of phage-bacterial interaction are countermeasures: phage-or plasmid-borne antirestriction functions [14–19]. In this regard, we note that *in vivo*, Iida and coworkers found the StySA restriction activity to be sensitive to the antirestriction activity of EcoP1 DarA [20], an activity otherwise only know to affect Type I restriction. Extension of Dar susceptibility to non-Type I enzymes would shed light on the scope of protection afforded by this phage countermeasure.

These defense mechanisms often cluster in genome islands. Genome comparisons between close prokaryote relatives led to the definition of variable “genome islands”, which are clusters of genes in one genome with no counterpart in another [6, 8, 21]. These variable genes usually deal with changing environments e.g. for resource acquisition and biological interactions. Some genome islands are specialized “defense islands”, enriched in genes that specifically regulate DNA entry [8, 22–26]. Defenses include RM systems (including modification-dependent nucleases, MDRS), and a wide variety of “abortive infection” elements. Abortive infection (Abi) processes prevent phage transmission to sister cells via cell death [27]. Abi mechanisms mostly target later events in phage development; even virion expression can occur (e.g. [28]). Defense islands may carry elements related to the Pgl (phage growth limitation) system in *Streptomyces coelicolor*, studied by M.C. Smith and coworkers [29–31], in which PglX-mediated methylation conditions the phage to restriction by MDRS upon infection of sibling cells. In this system the host genome is *not* methylated.

Sorek and co-workers extended the suite of methylation-protected defense systems using neighborhood analysis anchored by homologs of a component of the Pgl system, PglZ [32, 33]. This identified a set of systems designated BREX (<u>B</u>acteriophage <u>Ex</u>clusion), gene clusters of 4 to 8 genes, depending on the subtype. A BREX system from *Bacillus cereus* was studied experimentally in *B. subtilis* by Goldfarb et al.[32]. Further experimental characterizations include a system in *E. coli* strain HS [16, 34]; one in *Lactobacillus casei* [35]; and a plasmid-borne system from *Escherichia fergusoni* [36]. Though some elements of BREX are related to Pgl, the two families displayed important differences in biological endpoints, particularly the role of methylation, which protects BREX hosts [34] but elicits restriction by Pgl sibling cells.

SenLT2II (StySA) is one of three RM systems in *S. enterica* sv Typhimurium LT2. The other two are multicomponent ATP-dependent systems of Type I (SenLT2I; LT, StyLT in the early literature) and Type III (SenLT2III; SB, StySB). SenLT2II (StySA) was shown to carry out modification of adenine residues [37]. Its distribution may be limited to serovar Typhimurium; a survey of 85 other *Salmonella* serovars that included 11 testable for StySA found no SenLT2II (StySA) restriction activity outside of serovar Typhimurium [38]. For clarity we use the terminology from the restriction literature here: StySA, which is SenLT2II in the reference LT2 sequence.

We identified the genomic location of the StySA system while analyzing the sequences of *S*. Typhimurium-*S*. Abony hybrid strains[39]. The triply restriction-deficient laboratory shuttle host LB5000 was constructed to be a restriction-deficient but modification-proficient recipient for molecular genetic manipulations [40]. LB5000 (our STK005) carries a mutated allele of StySA (SenLT2II) named *stySA29*. This allele is also carried by our experimental host ER3625, descended from LB5000 [39]. LT2 locus_ID STM4495 is located genetically in the appropriate place to specify StySA. The single large protein specified by STM4495 was initially predicted to carry both R and M activities (REBASE 2015; 1225 aa; Type IIGS gamma). This is quite distinct from Type I enzymes, which require two genes for M activity and a third for R activity. Notably, this locus_ID is within a variable chromosomal island anchored by *leuX* where numerous non-homologous mobile elements are found in *E. coli* and *Salmonella* [41].

In this study, we describe the relationship of the StySA RM system to BREX systems and report the genetic contributions of its constituent genes to host protection and interruption of phage infection. The transcriptional organization of the locus contributes to understanding of relative expression levels in the native context. We find that *brxC* and *pglZ* mutations in our LB5000 descendant are responsible for the lack of restriction and modification; only the N-terminal domain of BrxC is required for modification. Serendipitously, we present evidence that the BrxL C-terminal domain by itself can have a large effect on host gene expression, particularly a non-SOS effect on resident prophages.

## Results

### Strain engineering design: domains, mutation clusters and transcription start sites

#### StySA is a BREX homolog

The non-restricting strain LB5000 and our experimental host ER3625 carried LT2-like m6A modification patterns, with one surprising exception: no modification attributable to STM4495 was found [39] despite identity of locus_ID JJB80_22595 to the LT2 sequence. Flanking genes corresponding to STM4492 and STM4496 did vary from the LT2 sequence (see further below; for a guide to terminology, see Materials and Methods).

Automated annotation [42] predicts a relationship of the StySA region with BREX and Pgl systems; we have adopted the automated name assignments from NCBI. The BREX homology is compelling when aligning the StySA region of LT2 (from [43])) with that of the *E. coli* HS BREX locus [34] (Figure 1A). In addition to the 6 genes described for the *E. coli* and *B. subtilis* BREX type I gene clusters, the LT2 StySA/BREX region carries an extra pair of genes, STM4493 and STM4494 (STM4493-4). STM4493-4 interrupt the similarity of the gene cluster (in the alignment gap Figure 1A). STM4493-4, which we will refer to as “accessory genes”, were apparently acquired by *Salmonella* separately from the genes we will designate “BREX homologs” (see Fig. 1 of [41]). We find below that neither of the accessory genes contributes to modification activity *in vivo* (see below “Phenotypic consequences of strain engineering”).

**Figure 1:**
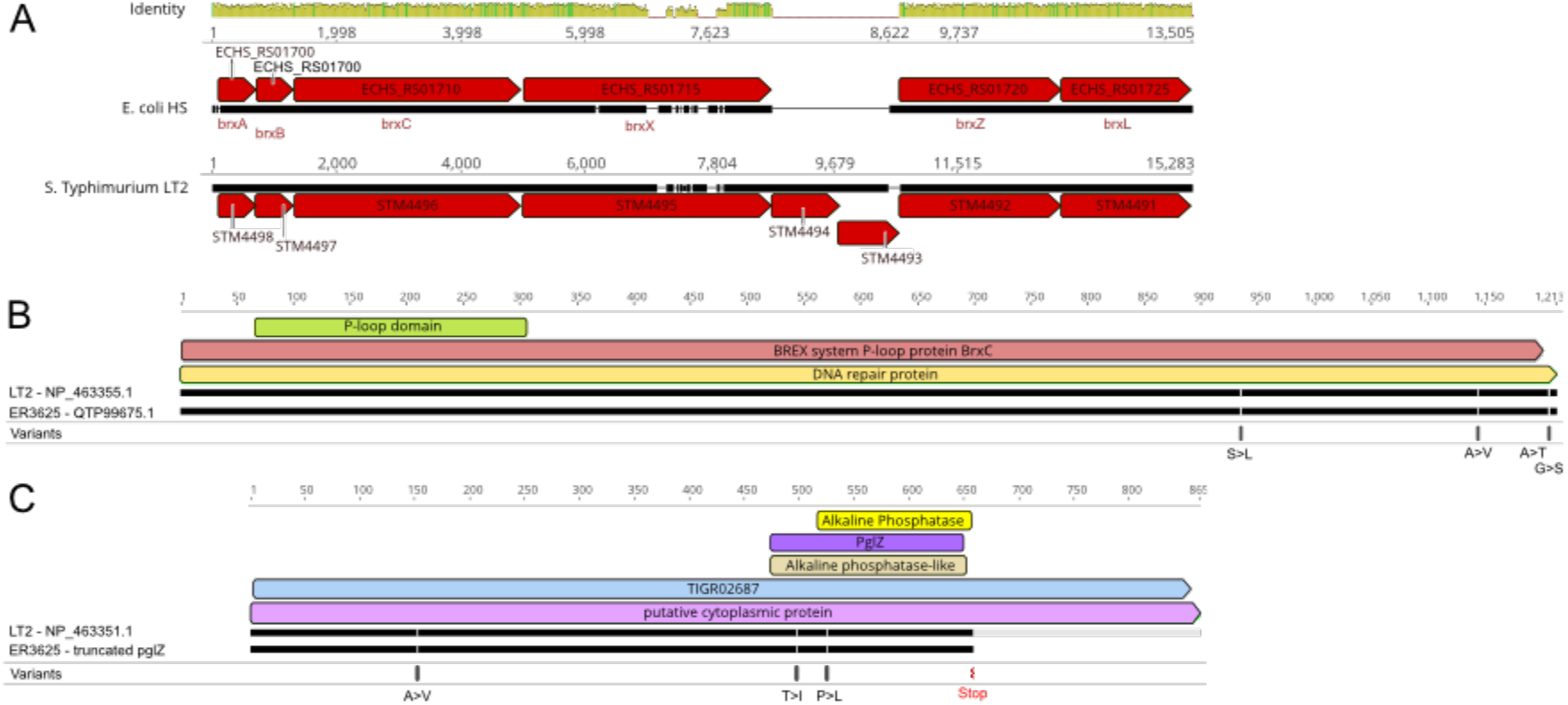
StySA locus similarity to *E. coli* BREX and protein variants in ER3625. Panel A, DNA alignment of BREX regions from LT2 with *E. coli* HS. Identity: green 100%, yellow close similarity, white sequence interruption. Numbers are nucleotide coordinates of segments extracted from the genome sequences (HS, LT2). Red boxes are annotated CDS with locus_ID numbers; black segments below the arrows represent DNA, with breaks where the two don’t match. A fine point is the small gap in the LT2 black line just upstream of *brxZ*: STM4491 and ECHS_RS01720. The black line for E. coli has a bit of sequence that has no counterpart in LT2. Presumably in E. coli there is a promoter/regulatory region for *brxZ*. For LT2 this function may be provided by 3’ end of STM4493 DNA. Panels B (BrxC) and C (PglZ) display amino acid coordinates for proteins of LT2 and its descendant ER3625. Breaks in the black lines between signify variant amino acid positions; the grey extension for the LT2 PglZ represents the C-terminal region missing in ER3625 (i.e. after the point of the translation termination for *pglZμ*). Colored boxes show predicted domains annotated (see text). Variant amino acids resulting from mutations present in ER3625 as compared to LT2 are indicated in the bottom rows of B and C.

#### Mutation set leading to StySA R^-^M^-^ phenotype for *stySA29*

ER3625 genes descended from STM4492 and STM4496 will be designated *brxCμ* (JJB80_22600, coding for a mutated version of BrxC) and *pglZμ* (JJB80_22580, coding for a mutated version of PglZ) respectively. Each carries multiple mutations relative to the LT2 ancestor. The amino-acid changes resulting from these are displayed in Figure 1B (BrxC) and Figure 1C (PglZ). The multiplicity of changes we attribute to the use of nitrosoguanidine mutagenesis during isolation of the *stySA29* allele [39, 44]. The R^-^M^+^ phenotype originally reported for the lineage [40] was unstable, losing modification (M^+^) ability [45]. Since tandem duplications are frequent in *Salmonella* [46], the original strain might have carried a duplication of the region, with clustered mutations in different genes in each copy, followed by resolution to single copy with both genes mutated and an R^-^M^-^ phenotype. Our genetic analysis below (Phenotypic consequences of strain engineering) is compatible with this scenario.

#### Genetic approach to the StySA locus

We chose to investigate this system *in situ*, replacing the mutated *pglZ* and *brxC* genes with wild type (WT). The engineering strategy first required a sketch of possible transcription signals, due to the complex organization of the locus and multiple potential toxic interactions. The strategy for gene deletion and replacement employed a method that leaves no scars, via an intermediate carrying a drug resistance cassette with its own promoter (see Materials and Methods and S2 Fig.). Strains created are listed in S2 File.

#### Transcription overview: StySA operon structure with Cappable-Seq and RNAseq

We first investigated transcription start sites with Cappable-seq [47] in the ancestral LT2 strain. The goal of such experiments is to better comprehend the complex organization of the BREX locus and the potential toxic interactions. It will also facilitate the design of deletion strains (see below). In our lineage with the reconstructed wild type StySA locus (JZ_058), we assessed the landscape of transcription using RNA-seq.

With Cappable-seq, two biological replicates identified 15,650 and 15,145 unique TSS positions in the genome (Material and Methods) (above the TPM >= 1.0 and EnrichRatio >= 2.5). Clustering was used to regroup close TSS giving a final count of 9,422 and 9,777 TSS clusters. Of these, 8,041 are shared positions with an adjusted R2 = 0.96426 and a P-value < 2.2E^-16^.

With this highly accurate method for mapping TSS location genome-wide, we confirm in our LT2 isolate (STK013) some TSS already made publicly available [48] and identify some new intragenic TSS (Figure 2). Primary TSS upstream of STM4498 (*brxA*) and STM4492 (*brxZ*) are confirmed; internal TSS are found within the STM4495 (*pglX*) and STM4494 (DUF4495) CDSs. A strong TSS cluster is located upstream of STM4490 (*mrr2*) in the forward orientation; this is adjacent to, but outside of the BREX/StySA cluster, defined as STM4491 (*brxL*)-STM4498 (*brxA*).

**Figure 2:**
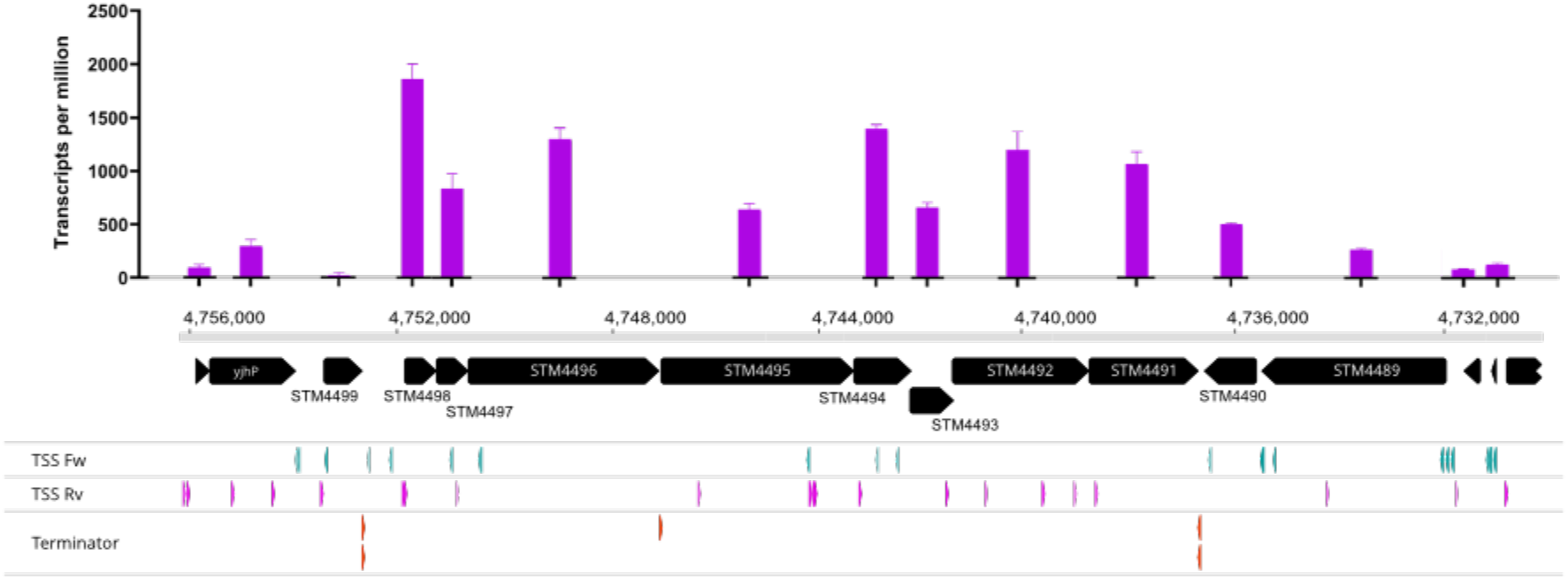
StySA locus operon structure analysis. Top panel corresponds to transcripts per million for the WT StySA (JZ_058) strain. Abcissa coordinates refer to LT2 sequence NC_003197.2. Black arrows are CDS (to scale), labelled with the locus_ID of LT2. The BREX-related genes are STM4498-4491; on the flanks, three (left) and five (right) external CDS are included to enable estimation of transcription reading into or out of the locus. The bottom panel show transcription start sites (TSS) determined in LT2 wild type using Cappable-seq. “Fw” is forward with respect to the genome coordinates, not the reading frames; “Rv” reverse. Transcription terminators were predicted using TransTerm.

RNASeq experiments allow us to map transcript profile and abundance (TPM) across the StySA locus in JZ_058, which has restored the locus to sequence identity with LT2 in the ER3625 lineage. We observe agreement between transcript abundance for each feature and available TSS for the ancestral LT2 region in STK013: the RNA profiles along the locus show varying transcript levels with alternating regions of higher and lower abundance. We infer likely transcription starts directly upstream of STM4498 (*brxA*), and within STM4494 (DUF4495) and STM4492 (*pglZ*) homologs. For the TSS experiment, we observed the TSS in front of STM4496 (*brxC*) was weaker than that in front of STM4498 (*brxA*); the TPM data suggest that this *brxC* promoter could be as strong as the others in this strain background.

Rho-independent terminators are predicted at three positions in this locus. Two are bidirectional, flanking the StySA locus (before STM4498 (*brxA*) and after STM4491 (*brxL))*. The locus should thus be transcriptionally insulated (see “Local and global inferences from transcriptome profiling” below).

The third predicted terminator lies between STM4496 (*brxC*) and STM4495 (*pglX*), suggesting multiple operons within the locus. Experimental evidence for the *brxC* terminator was sought, using a combination of SMRT-Cappable-seq and RACE methods (S9 File). The combination approach (S9 panel A) supports the authenticity of the *brxC* transcription terminator: read number drops at the end of *brxC* (S9 panels B and C), with some read-through to *brxX*. Coverage was too low for stronger conclusions (~15 reads before the terminator position and ~5 after it).

These experiments suggest that STM4494, STM4493, *pglZ* and most likely *brxL* are on one transcript (S9 panel D), while *brxA, B* and *C* and *pglX* are on a different transcript (S9 panel A).

#### Putative functions of the BREX core gene components and response to genome manipulation

For each annotated CDS in the ER3625 StySA locus, gene segments were deleted with consideration of the transcript structures predicted from the analysis above. The engineering strategy for gene deletion and replacement employed a method that leaves no scars, via an intermediate carrying a drug resistance cassette with its own promoter (see Materials and Methods and S2 Fig.). Strains created are listed in S2 File

Functional domain annotations were described for the Pgl and BREX systems. The characterized Pgl system expresses a putative phosphatase (*pglZ*), a protein kinase (*pglW)*, a candidate adenine-specific DNA methyltransferase (*pglX*) and a P-loop ATPase (*pglY*) [29]. Proteins specified by two of the 6 genes in type I BREX are similar: PglZ and PglX. BREX-specific genes specify BrxL, identified as a Lon-like protease-domain; BrxA, proposed to be a structural homologue of RNA-binding antitermination protein NusB; BrxB, a protein of unknown function and BrxC, a large protein with a P-loop ATP-binding domain [32]..

Schematic domain predictions are shown in Figure 1B (*brxC*, STM4496 in LT2; JJB80_22600 in ER3625), Figure 1C (*pglZ*, STM4492 in LT2; JJB80_22580 in ER3625) and in S1 File (*brxA;* STM4498 and JJB80_22610; *brxB*, STM4497 and JJB80_22605; *pglX*, STM4495 and JJB80_22595; STM4494 (JJB80_22590), STM4493 (JJB80_22585) and *brxL*, STM4491 and JJB80_22575).

From left to right in Figure 1A, STM4498 codes for a BrxA homolog. The gene contains a putative RBS for STM4497. Domain prediction of DUF1819, is inferred to code for an inner membrane protein. We were unable to create a *cat* replacement of this gene.

STM4497 codes for a BrxB homolog, annotated as a DUF1788 protein. Replacement of *brxB* with *cat* has extensive effects on transcription of the cluster. The *cat* replacement is viable in three allelic states (see “Local and global inferences from transcriptome profiling” below).

STM4496 codes for a BrxC/PglY homolog, originally annotated in LT2 as “putative ATPase involved in DNA repair”. Geneious implementation of InterProScan with LT2 protein sequence AAL23314 (NP_463355.1) yielded domain hits including an N-terminal ATPase region (P-loop NTPase, ATPase involved in DNA repair, P-loop containing nucleoside triphosphate hydrolase) and a C terminal region designated “SMC N-terminal domain”. In ER3625, the four mutations in *brxCμ* (JJB80_22600) are clustered in the 3’ end of the gene (Figure 1B). InterProScan detects only the NTPase hits in the N-terminus in the corresponding protein (QTP99675.1). Presumably the variant amino acids result in loss of the SMC-domain recognition.

STM4495 is a BrxX/PglX homologous gene, with signatures of m6A methyltransferases. Domain and annotation search leads to N-6 adenine-specific DNA methylases hits. DNA alignment with the *E. coli* HS homolog show two patches of divergence (Figure 1A). This is compatible with potential DNA recognition in that region, since the two MTase recognize and modify different motifs. Methyltransferases often segregate site specificity into a recognition module [49–53]. We were unable to make a deletion of this gene in three different allelic states. Toxicity due to disruption of transcription in the intermediate or a toxic protein complex formed from other components in its absence are possible explanations.

STM4492 is a BrxZ/PglZ homolog, thus putatively has phosphatase activity. PglZ of *Streptomyces coelicolor* was required for viability in the presence of PglX; neither gene deletion nor point mutation of the candidate catalytic aspartates in the phosphatase fold could be made in single copy [29]. Geneious implementation of InterProScan with LT2 protein sequence AAL23310 (NP_463351.1) yielded hits with four different alkaline-phosphatase-like domains; these are found in alkaline phosphatases, arylsulphatases, phosphoglycerate mutase, phosphonoacetate hydrolases or phosphoenolmutases.

The mutated homolog in ER3625 (*pglZμ;* JJB80_22580) carries five mutations. Notable is a stop codon just 3’ to the annotated PglZ domain. The N-terminal protein fragment (here designated PglZ’) would carry three variant amino acids when translated; a fifth mutation is distal to the stop. No protein corresponding to PglZ’ is listed in the NCBI proteome for ER3625, although the gene is annotated. To determine whether PglZ’ is recognized by domain-detection programs, a translation of the annotated gene (truncated at the stop) was submitted to InterProScan. The PglZ annotation was returned to this search (Figure 1C); thus the structural domain PglZ/AlkPhos is recognized even when the tail of the protein is not there. This does not prove functionality but suggests that partial assembly or partial enzymatic function could be retained. The intact gene is required for both R and M activity *in vivo* (see below). In addition, we have strong evidence that the stop codon is polar on downstream transcription.

STM4491 (JJB80_22575) is a BrxL homolog. Domain predictions for protein sequence AAL23309 (NP_463350.1) gave multiple hits to Lon-protease domains. This is proposed to be two-domain Lon-like protease structure with an N-terminal ATPase and a C-terminal proteolytic domain. However, it is most likely not a protease as the catalytic site is not present (see further below).

### Phenotypic consequences of genome engineering

Five phenotypic measurements were carried out with the strains constructed: phage restriction; modification at StySA sites (using Dam modification as a control); growth rate; expression level of the M gene (*pglX*, STM4495; JJB80_22595); and for a selection of strains, RNAseq was used to analyze local and global transcription effects.

Colony formation was found to be slow and variable on plates for some strains and not others (See S3 Fig.). Thus, a comparison of growth rate in liquid was undertaken.

#### PglZ and BrxC variants contribute to both R^-^ and M^-^ properties

Only one R^+^ strain was obtained. JZ_058 was able to restrict phage L; it carries wild-type LT2 sequence in place of both *brxC* and *pglZ* of ER3625. The magnitude of restriction agreed with literature reports [40, 54]: 100-fold reduction in plaque-forming ability (S3 Fig.).

Modification *in vivo* did not require wild-type BrxC: when *pglZ* is wild-type, mutated *brxCμ* allows modification (Figure 3 panel B, JZ_022). However, the N-terminus of BrxC is required for modification, since Δ*brxC::cat* no longer modifies (Figure 3 panel B, JZ_040). In contrast, modification required PglZ to be intact (Figure 3 panel C: JZ_028 and JZ_043).

**Figure 3:**
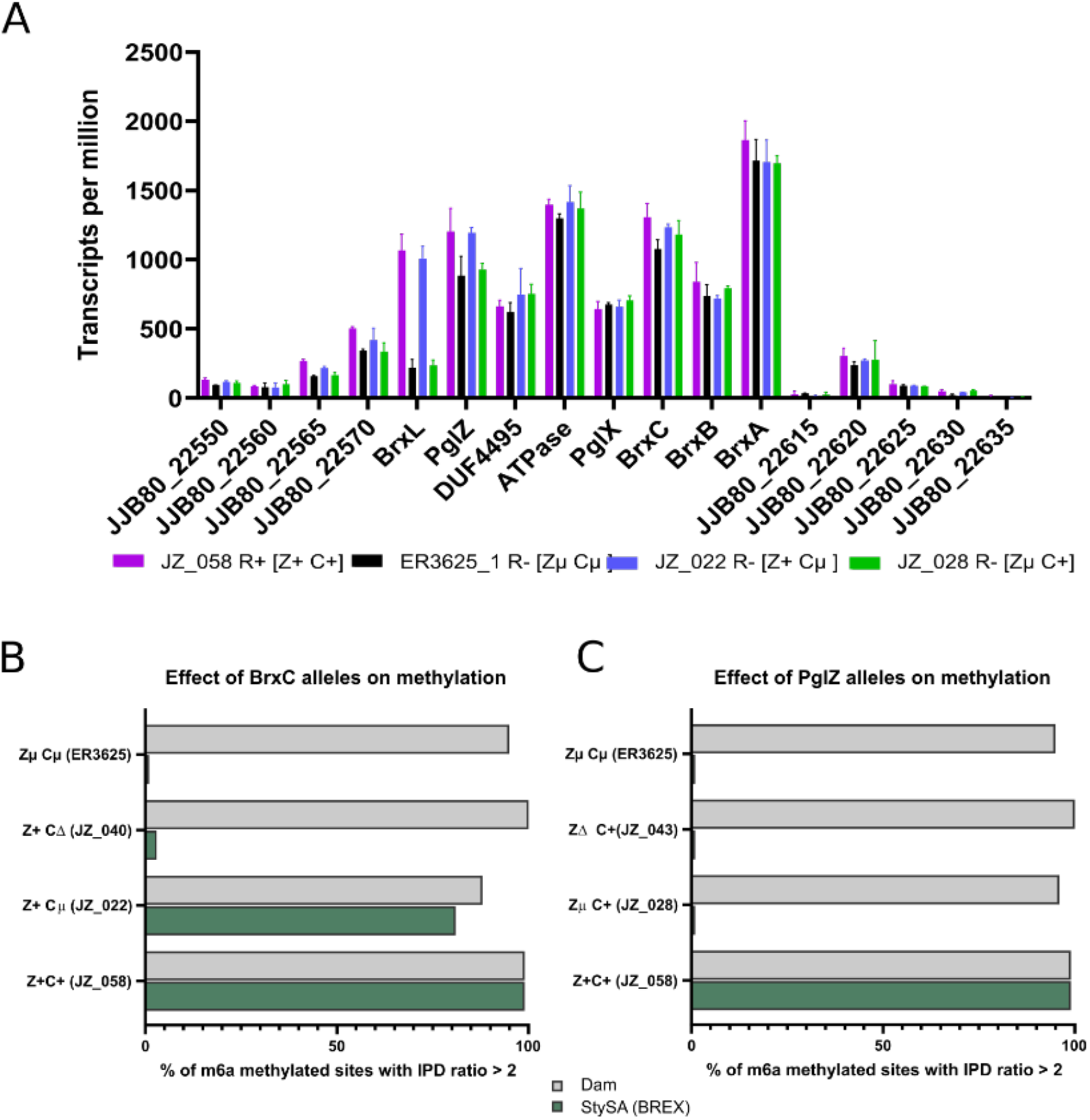
BrxC and PglZ alleles determine restriction and methylation phenotypes, and BrxL transcript abundance. Panel A: Transcription level across features in the StySA gene neighborhood for four strains with clustered mutations (μ) or wild type (+) in *brxC* and *pglZ*. Strain restriction phenotypes are R^+^: 100-fold reduction in plaque formation with phage L; R^-^ no restriction activity. Features shown (in opposite orientation to Figures 1 and 2; not to scale) are NCBI-assigned locus_ID (outside the StySA cluster) or gene names (within it); vertical axis is transcripts assigned per million transcripts sequenced. *pglZμ* is polar on *brxL*. Panels B and C: effects of *brxC* (Panel B) and *pglZ* (Panel C) alleles on methylation of StySA sites (green bars) with Dam sites (gray bars) as a control.

#### Methylation activity does not depend on the level of PglX (MTase) expression

We used qPCR to measure *pglX* transcription levels to clarify the question of whether the degree of methylation is regulated by changes in transcription that result from engineering steps. The *pglX* transcript level does not correlate with M phenotype (**Error! Reference source not found**. panel B top, M^+^, StySA modified; M^-^ StySA unmodified; M^p^, partial modification). Significant differences in transcription are seen in some strains relative to ancestor ER3625, but the highest transcription is found in unmodified strains. A negative control is OD_127, a strain with a multigene deletion removing *pglZ-brxC* and including *pglX*.

#### Methylation activity is not affected by ΔSTM4493::*cat*, ΔSTM4494::*cat* or Δ*brxL::cat*

Removal of accessory genes STM4493 or STM4494 do not affect methylation (Figure 4 panel A top). M^+^*pglZ^+^ (brxCμ* or *brxC^+^*) hosts retain modification with ΔSTM4493::*cat* (JZ_091 or JZ_080) or ΔSTM4494::*cat* (JZ_092 or JZ_069); modification did not return to the M^-^ *pglZμ brxC^+^* host (JZ_094 and JZ_095). Similarly, removal of the N-terminal segment of *brxL* did not change M status (Figure 4 panel A bottom). The contradictory properties of Δ*B::cat* are discussed further below.

**Figure 4:**
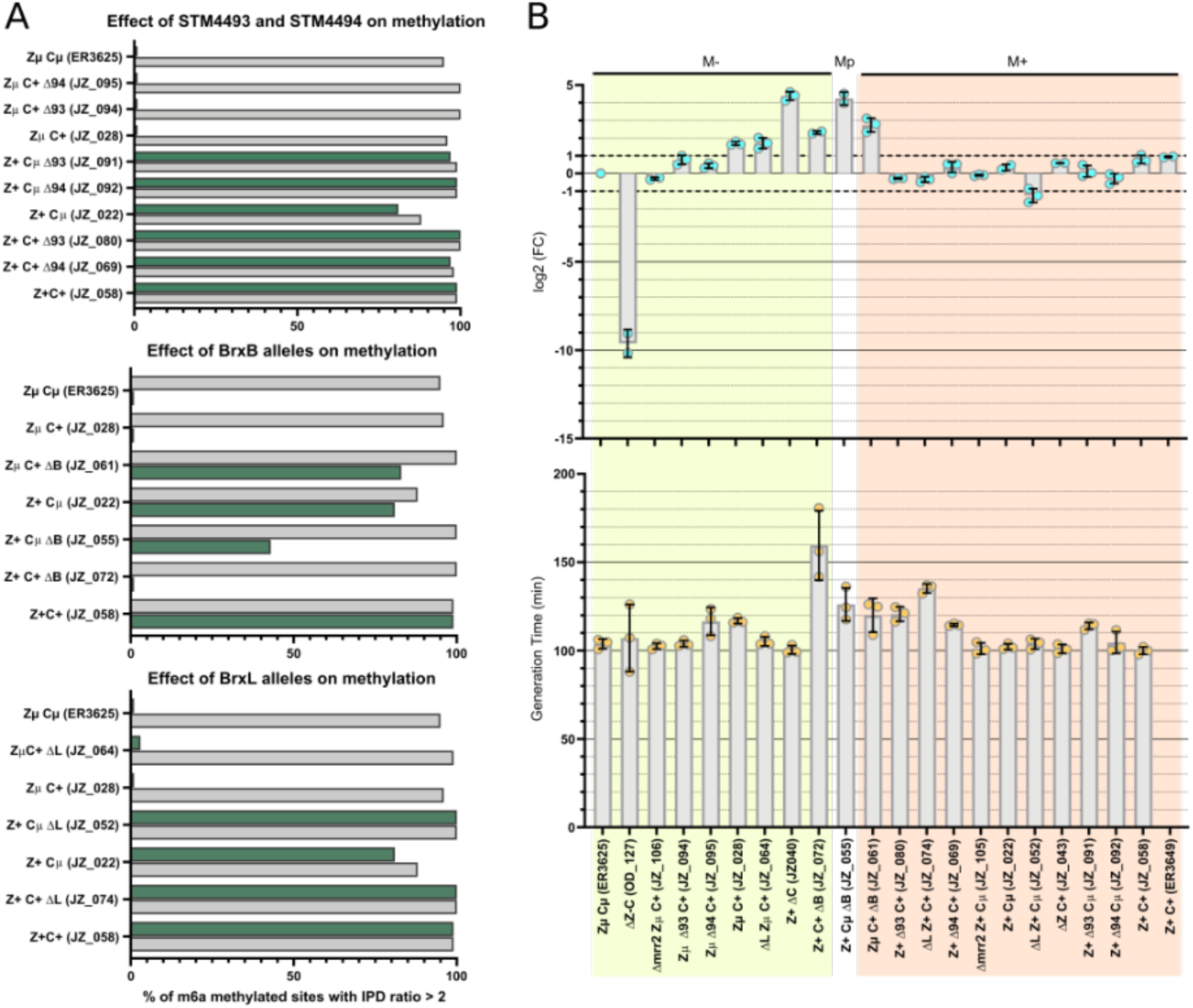
Modification, PglX transcript and growth rate responses to engineering. Panel A, Methylation of Dam sites (gray bar, internal control) and StySA sites (green bars) with different genetic configurations. μ, mutation cluster inherited from LB5000; + wild type LT2 replacement; Δ, segment of the gene replaced with a *cam* cassette; (), strain number. Panel B, STM4495 (*pglX*) expression in the different strains. Each dot represents a biological replicate, the bar chart represents the mean relative expression level. The methylation level is simplified and represented by M-, Mp and M+ respectively for methylation < 5%, methylation at an intermediate level and methylation > 90%. Panel C, Generation time in minutes. Each dot represents a biological replicate, the bar chart represents the mean relative expression level.

#### Growth rate is affected by Δ::*cat* constructions

We found on plates that the strains have different colony phenotypes and different growth rates (See S3 Fig.) so we did a liquid culture experiment in plates to accurately compare the strains growth, see Figure 4 panel B bottom. Most strains show growth rates about the same as the ancestor ER3625. Interestingly, there is no correlation between methylation level and loss of fitness. However, the strains with Δ*brxB::cat* (JZ_072, JZ_055 and JZ_061) share slower growth rate, regardless of the allelic state of *brxC* and *pglZ*. We attributed this shared property to the unbalanced expression of the whole operon due to the *cat* cassette promotor, see below and Figure 5. Of the three Δ*brxL::cat* strains, the one closest to wildtype also grows more slowly: JZ_074 (Δ*brxL::cat pglZ^+^ brxC^+^)*.

**Figure 5:**
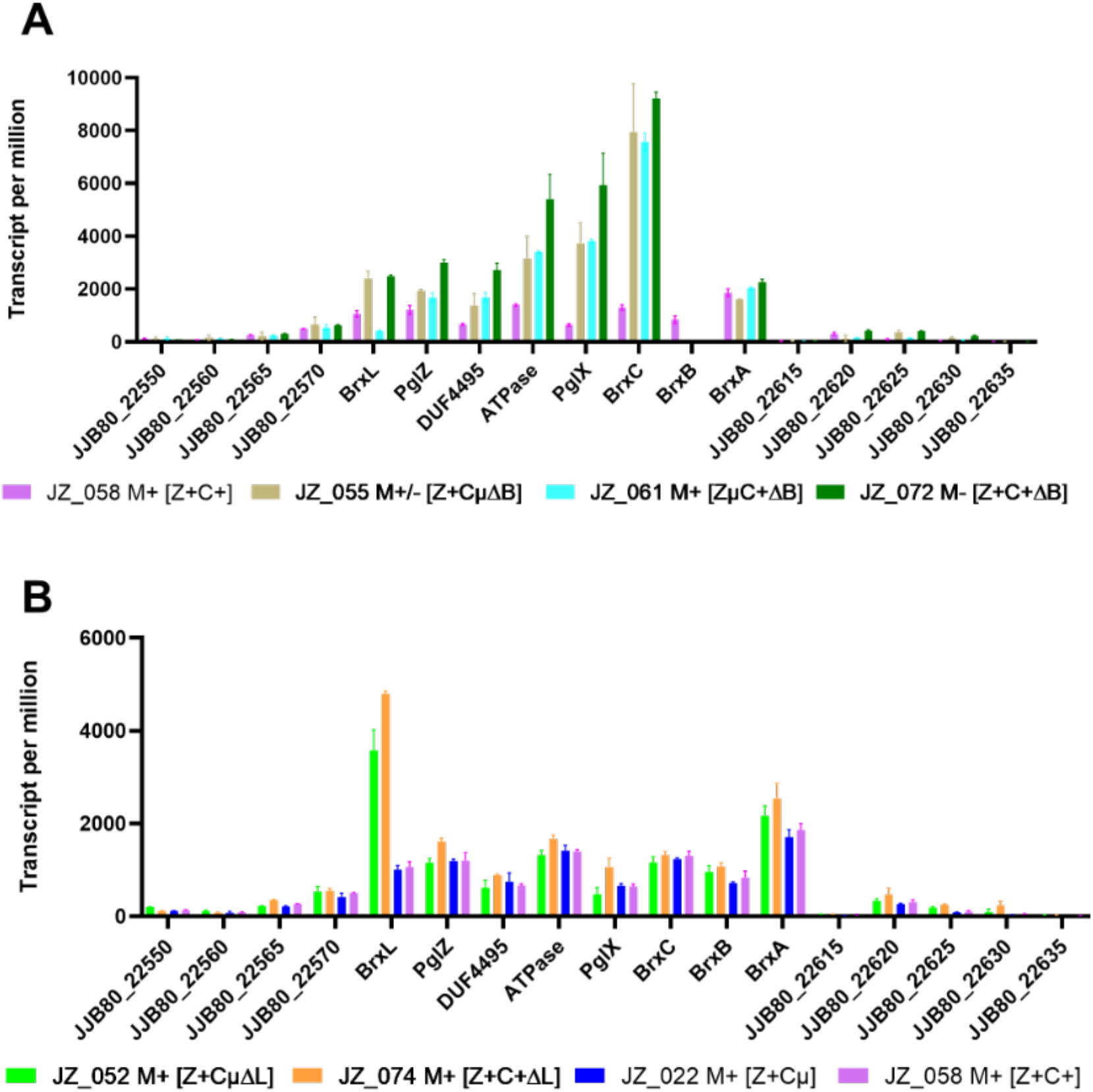
StySA locus transcription with Δ*brxC::cat* and Δ*brxL::cat*. The abcissa displays features in the chromosome region with StySA/BREX locus and flanking features, not to scale, in order of chromosomal coordinate. Transcription from *cat* and all CDSs within StySA is right to left (opposite to Figures 1 and 2). Ordinate is transcripts mapping to the feature per million. Panel A: Strains: brown, *Z^+^Cμ* Δ*brxB::cat;* blue, *ZμC^+^* Δ*brxB::cat;* green, *Z^+^C^+^* Δ*brxB::cat;* violet, *Z^+^C^+^* (wild type). A lower plateau covering *pglX* and ATPase (STM4494) for each strain following *brxC* may result from action of the *brxC*-distal terminator. A similar lower plateau covering DUF4495 (STM4493) and *pglZ* suggests additional termination following ATPase, for unknown reason. Lower transcription of *brxL* in JZ_061 is consistent with polarity seen in Figure 3 panel A. Panel B: In Δ*brxL::cat*, the remnant 807 nt segment of *brxL* shows very high transcription from the *cat* promoter in Δ*brxL::cat* relative to isogenic *brxL^+^*. This transcription does not leak through the bidirectional terminator into flanking features, e.g. JJB80_22570. Strain pairs: *Z^+^Cμ* background--green Δ*brxL::cat*, blue *brxL^+^; Z^+^C^+^* background--orange *ΔbrxL::cat*, violet *brxL^+^*. There may also be a small increase in transcription of *brxA-brxB* in the *ΔbrxL::cat* strains.

### Local and global inferences from transcriptome profiling

#### *pglZμ* is polar on *brxL*

RNASeq measurements of transcription of the StySA locus (Figure 3 panel A) suggest that the alternative alleles (μ or WT) do not affect transcription within the BREX cassette, with one exception: transcription of *brxL* is significantly increased when *pglZ* is wild type (JZ_058 and JZ_022) relative to *pglZμ* (ER3625 and JZ_028). We infer that translation termination at the *pglZ* stop codon is polar on transcription of *brxL*. The phenomenon of translational polarity is well known though the mechanism is still under study (e.g., [55, 56]); transcript quantity is decreased following a translation stop. However, the R^-^ phenotype is not reversed simply by relief of polarity and increased BrxL expression; JZ_022 carrying *pglZ^+^ brxCμ* does not restrict (S3 Fig.). Intact BrxC is also needed.

#### Overexpression of ‘*brx*L yields global effects without leakage across the insulating terminator

RNAseq results for Δ*brxL::cat* strains provide strong evidence that the left-hand terminator insulates the flanking sequence from readthrough (Figure 5B). Inside the locus, only the remnant ‘*brxL* gene has altered expression; this is strongly increased relative to isogenic *brxL^+^*.

There are other effects of this disruption design. Growth rate is impaired for Δ*brxL::cat* JZ_074 (*brxC^+^*) though not JZ_052 (*brxCμ*) (Figure 4B bottom panel); for both strains, structural genes for two prophages are overexpressed, as well as numerous other genes outside the locus (Figure 6, S5 File, S10 File).

**Figure 6:**
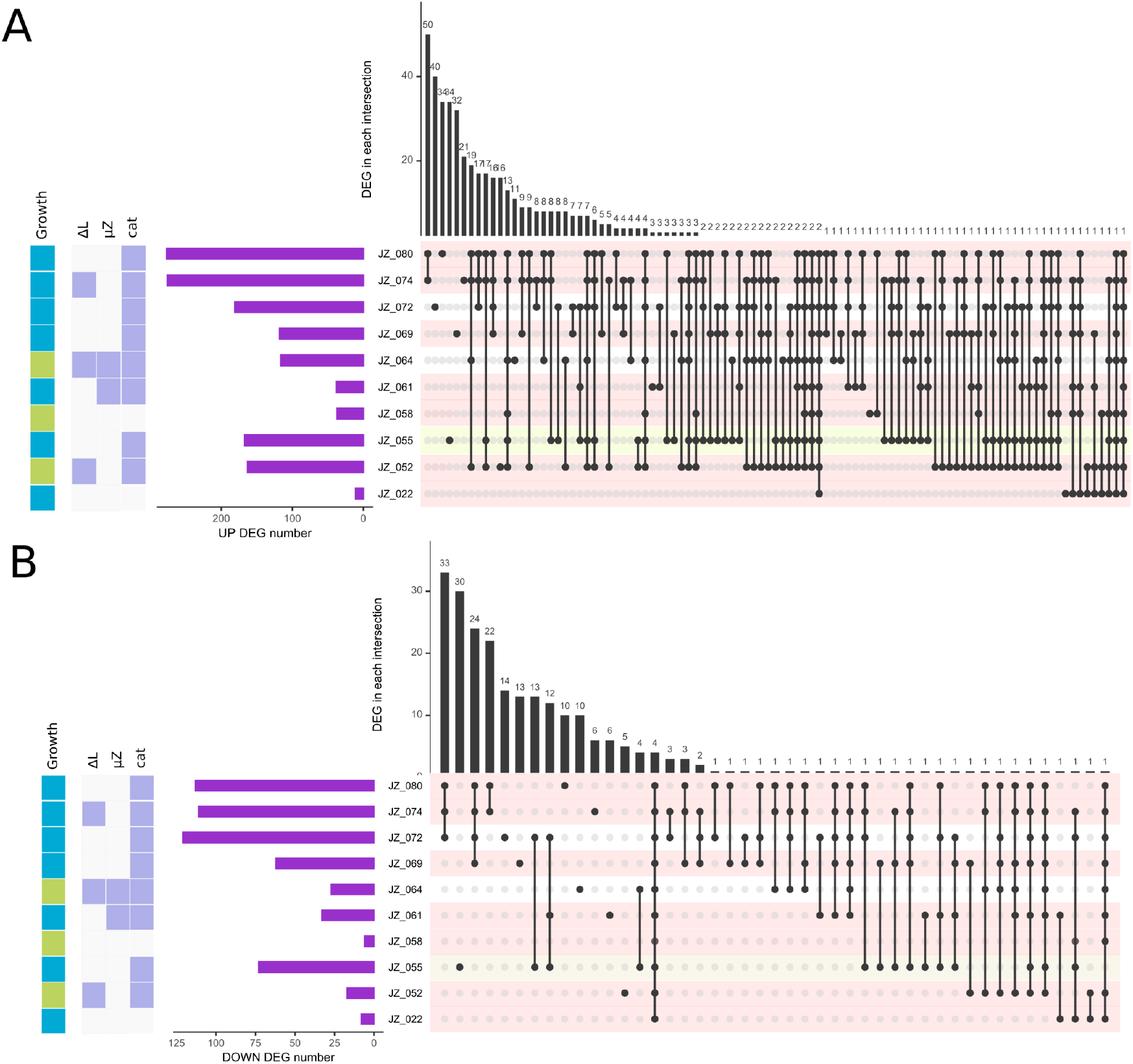
Global changes in transcription visualized with UpsetR. Transcription for each CDS feature in each strain was compared with ER3625, with differentially expressed genes (DEG) reported as adjusted pvalue. On the right, sets of genes with changed expression and the sharing of sets with other strains are represented by dots connected by lines. In each row, a black dot represents a set of genes in a particular strain. Between rows, lines connect that dot (set) to other strains with dots representing the same set. The histogram at the top indicates the number of genes represented in that set (intersecting group). Top and bottom panels are respectively upregulated and downregulated sets of genes. In each row, levels of methylation are coded pink (M+), yellow (Mpartial) and white (M-). On the left, the purple bar chart indicates the total number of DEG in each strain. In the far left column, each strain was assigned a growth property, blue for slow growth, green for normal growth. In the middle columns, selected strain genotype information is color coded: white, marker is absent; blue, marker is present. Markers are: ΔL, the *cat* cassette has replaced the 5’ segment of *brxL;* μZ, multiple mutations are present in *brxZ;* cat, the chloramphenicol cassette and its promoter are present. JZ_028 (*brxC^+^ pglZμ*) does not appear in the figure because it shows no change from ER3625 in phenotype or gene expression. The *cat* cassette is present in *brxB*: JZ_055, JZ_061, JZ_072; STM4493 (DUF4495): JZ_080 and in STM4494 (ATPase): JZ_069.

#### Unbalanced expression of BREX locus components with Δ*brxB::cat*

The modification properties of strains with deletion/replacement of *brxB* are contradictory. In the genetic context of unmethylated strain *brxC^+^ pglZμ* (JZ_028), the *cat* replacement of *brxB* (JZ_061) restores methylation (Figure 4 panel A, middle graph). In contrast, with modified strain *brxCμ pglZ^+^* (JZ_022), *cat* replacement of *brxB* in JZ_055 leads to a partial loss of methylation. In wild type *brxC^+^ pglZ^+^*(JZ_058) it leads to complete loss of methylation (JZ_072). In Discussion, we suggest that component ratios may account for this.

The “partially methylated” phenotype was reproducibly found in the genotype *brxCμ pglZ^+^* Δ*brxB::cat*. Three independent constructions (JZ_055, JZ_056 and JZ_057) with the same configuration all display the same modification pattern. Dam sites are methylated normally. Examining the distribution of modified StySA sites among these isogenic strains, we observed in each a patchy modification pattern. Multiple nearby StySA sites have very low modification while others comprise fully-modified regions, rather than consistent half-modification (S6 File).

These RNAseq results for Δ*brxB::cat* strains also support effective action of the terminator between *brxC* and *pglX*: the transcript drops by half between these genes (Figure 5A). This is consistent with the 5’RACE evidence for effective transcription termination at the end of *brxC* (see above “Transcription overview” and S9 File). Termination is not complete, allowing readthrough to *pglX*.

#### Global inferences from transcriptome differential expression profiling

The three engineered strains lacking *cat* cassettes are least globally affected in terms of gene expression (Figure 6). Interestingly, JZ_028 (*brxC^+^ pglZμ*) with unchanged R^-^M^-^ phenotype, has no significant differences with the ancestor (and does not appear in Figure 6). JZ_022 (*brxCμ pglZ^+^*) has recovered modification (M^+^); outside the locus, 11 genes show increased expression, 8 decreased. JZ_058 (*brxC^+^ pglZ^+^*) has recovered both M^+^ and restriction (R^+^) with 37 changes up, 6 changes down. Of the 11 genes with increased expression in M^+^ JZ_022, 2 are unique to it: a possible operon of genes JJB80_02810 (PLP-dependent aminotransferase) and JJB80_02815 (M15 family metallopeptidase). Of the 37 genes increased in R^+^M^+^ (JZ_058), 1 is unique: JJB80_02375, a ribosomal protein associated with changes in frameshifting. It seems unlikely that these extra-locus changes. are responsible for the M and R phenotypes.

A very striking result is that strains with *cat* cassettes have very large numbers of genes with significant changes in expression, and the sets of genes with altered expression are not shared (Figure 6). We think it unlikely that the chloramphenicol acetyltransferase enzyme itself is responsible; no drug was present. If these effects were due to the action of the Cat protein (chloramphenicol transacetylase) without substrate, all the *cat*-containing strains should have similar expression impact. However, both number of genes and the shared sets are quite different. For example, with *cat* cassette in *brxB*, the *pglZμ* strain JZ_061 has a fairly small number of genes affected--a number similar to the number changed when wild type (*brxC^+^ pglZ^+^*) is restored, with no cassette (JZ_058). The three Δ*brxL::cat* strains have a much bigger effect, which is even enhanced by the presence of WT alleles of Z and C.

Instead it is likely that unbalanced expression of elements of the StySA island mediate these drastic global effects. In most cases six, three or two downstream genes are overexpressed. Possible action on off-target substrates by helicase, phosphatase and ATPase activities is greatest for *brxB* (JZ_055, JZ_061, JZ_072) with overexpression of *brxC, pglX*, STM4493-DUF4495, STM4494-ATPase, *pglZ* and *brxL*. For STM4493-DUF4495 (JZ_080) downstream genes are STM4494-ATPase, *pglZ, brxL*. For STM4494-ATPase (JZ_069), only *pglZ* and *brxL* are overexpressed. In general flavor, these global changes affect the three prophages in the strain and cell surface composition (S5 File).

Global changes in the Δ*brxL::cat* strains JZ_064, JZ_072 and JZ_074 are more interesting, because the local effect of the *cat* insertion is limited: only the fragment of the *brxL* gene, *‘brxL*, is overexpressed (Figure 5B). Genes lying outside the StySA cluster are insulated from transcription emanating from within by a very strong terminator. The strong effects on global transcription (Figure 6 and S5 File) suggest that the *‘brxL* transcript or a translation product act outside the locus.

We favor the model that this partial gene is translated into a stand-alone protein embodying the C-terminal domain (Lon_C, SSF54211: “Ribosomal_S5_D2-typ_fold”; see S7 File). The 5’ 1278 nt segment of the gene has been replaced with the *cat* cassette, but the 3’ 807 nt still carries an ORF. Translation of the original *brxL* CDS terminates within the added *cat* cassette (118 nt + 10 nt cassette). A transcript from the *cat* promoter could allow translation restart following the *cat* CDS, at a CTG 20 nt into the *brxL* remnant (52 nt from the *cat* UGA). Such translation would yield a protein carrying the signature motifs from three domain annotation sources: GENE3d:3:30 Ribosomal_S5_D2-typ_fold_subgr; PFAM: Lon_C; Superfamily SSF54211 (Ribosomal_S5_D2-typ_fold). The candidate activities of the C-terminal domain include hydrolysis, phosphoryl transfer and folding, any of which could explain major cellular effects.

## Discussion

### Genetic states

Not all deletion derivatives designed were successfully constructed. Some reasons for this are addressed here. Failure to recover deletion designs can be interpreted two ways: overexpression of downstream genes in the cassette-containing intermediate strain may be toxic; or the target protein itself is required for protection from toxicity of other components of the locus.

Toxin/antitoxin pairs are frequently associated with defense islands [8]. Removal of the antitoxin alone is usually lethal in such cases, while removal of the toxin alone is tolerated. There are two possible such pairs within the StySA locus: BrxA-BrxB and STM4493-STM4494.

Our failure to construct *ΔbrxA* in three genetic contexts might be expected if BrxB were a toxin and BrxA an antitoxin. However secondary transcription effects are also anticipated (e.g. results of Δ*brxB::cat*), so the inference is weak.

#### STM4493-STM4494

We find that both genes can be disrupted individually in three allelic contexts (Figure 4; S2 File). This argues against action as a toxin/antitoxin pair, at least in this genetic environment.

We were also unable to recover a deletion of *pglX* in three allelic contexts, even with a parent strain already lacking modification. Potentially the intermediate would express STM4493-4494 aberrantly, rescuing a potential toxic effect not found otherwise. We have not evaluated potential expression of a truncated PglX protein.

### Components of methyltransferase

The N-terminal domain of BrxC and the complete PglZ protein are required for DNA modification (Figure 3 panels B and C), while the C-terminus of BrxC is also required for restriction (S3 Fig.). Modification and restriction are restored without notable effects on transcription within the locus, except for relief of polarity on *brxL* (Figure 3 panel A).

In agreement with findings for a related system from *E. coli* HS [34], BrxL is required for restriction but not methylation. Accessory elements STM4494 and STM4493 are not essential for methylation.

BrxB is also not required for methylation activity and is unlikely to participate in the active complex, though it could act as a chaperone or to support subcellular localization. For example, overproduction of one component of transposase can result in autoinhibition[57]; with multicomponent replication complexes, dominant-negative phenotypes can result from formation of unproductive complexes or failure to remodel them[58].

The contradictory effects of the Δ*brxB::cat* allele in the three genetic contexts (Figure 4 panel A middle) is likely to relate to the mutational status of the *brxC* and *pglZ* proteins overproduced (Figure 5A), rather than the expression level of StySA components per se. The effect on StySA-BREX locus transcription is similar for all three strains: transcription resulting from the strong *cat* promoter is 10-fold higher than in the wild-type state for *brxC*, declining to ~twice the wild type in the *brxL* gene. Once again, the effect on *brxL* transcription of the stop codon in *pglZμ* is seen.

### Overexpression of a BrxL fragment leads to action outside the StySA locus

The genome-wide transcription effects of Δ*brxL::cat* are seen in three genetic contexts, providing strong evidence for function of the truncated gene (Figure 6, S5 File). We propose that this allele provides local opportunity for translation of a truncated protein domain (S7 File). Such a solo domain might well escape control on its action normally mediated by association with the other proteins in the BREX cluster. (see “Global inferences from transcriptome profiling” and S5 File and S10 File). The intact BrxL protein is critical to restriction action against phage L here (S3 File), in agreement with [34]. A striking aspect of these global effects is enrichment for expression of prophage genes, particularly Fels-1 and Gifsy-1 (S5 File). Regulation of expression in these three prophages is tightly intertwined, mediated by antirepressor interactions [59] and related to expression of pathogenicity functions [60]. Interestingly, the replication regions of these phages are not overexpressed, while components of the virion are--the phages are not induced to replicate, as would occur with DNA-damage induced escape. This activity may shed light on the phage replication-inhibiting mode of action the role of the intact restricting complex BrxL.

## Material and methods

### Terminology

- StySA: genes and phenotypes related to the cluster of genes that specify REBASE system *SenLT2II* of organism number 18099 and its lineal descendants [39]. This cluster is characterized genetically in this work, using ER3625 as experimental system. This segment of the ER3625 genome is descended from LT2 with mutagenesis, although two distant genome segments of ER3625 descend from *Salmonella enterica* serovar Abony SW803 [39].
- RM: restriction-modification system
- R^+^, R^-^: restriction phenotype, measured by reduction in plaque formation of bacteriophage L when grown on a StySA M^-^ host [61]
- M^+^, M^-^, M^+/-^: StySA sites (GATCAG) are methylated at the second A fully, not at all or partially.
- *gene*::Δ*cat*: a gene with a portion deleted and replaced with the chloramphenicol resistance cassette of pKD3, including a strong promoter.
- *brxC*, BrxC: gene, protein specified by LT2 locus_ID STM4496
- *brxCμ*, BrxCμ: gene, protein specified by ER3625 locus_ID JJB80_ 22600
- *pglZ*, PglZ: gene, protein specified by LT2 locus_ID STM4492
- *pglZμ*, PglZμ: gene, protein specified by ER3625 locus_ID JJB80_ 22580

The rest of the DNA sequence of the ER3625 BREX cluster is identical in sequence to that of LT2, so the genes and proteins will sometimes be referred to using the LT2 nomenclature. This will allow more ready access to database information since the LT2 sequence is well curated.

- *brxA*: STM4498; JJB80_22610
- *brxB*: STM4497; JJB80_22605
- *pglX*: STM4495; JJB80_22595
- STM4494: JJB80_22590, ATPase
- STM4493: JJB80_22585, DUF4495
- *brxL*: STM4491; JJB80_22575
- ‘*brxL*, ‘BrxL: N-terminally truncated gene and potential product resulting from replacement of the 5’ end of *brxL* with the *cat* cassette of pKD3, as described in S7 File.

### Genome engineering

The Datsenko and Wanner method [62] was adapted and combined with the FAST-GE method [63] to engineer ER3625 descendants (Figure 7). During the second step of classic *λ* Red engineering, drug resistance cassettes would be removed by FLP/FRT recombination. For this, PCR-amplified *cat* cassette must be flanked by FRT (FLP Recombinase Target) sites such that the Flp recombinase can drive specific recombination between FRT sites. We found that resident FRT sites in ER3625 (*aroA::FRT, mrr::FRT::kan*) interfered with cassette addition to new sites, with the newly synthesized fragment recombining preferentially into the resident sites. Thus, in this work, the *cat* cassette was amplified and inserted without FRT sites.

**Figure 7:**
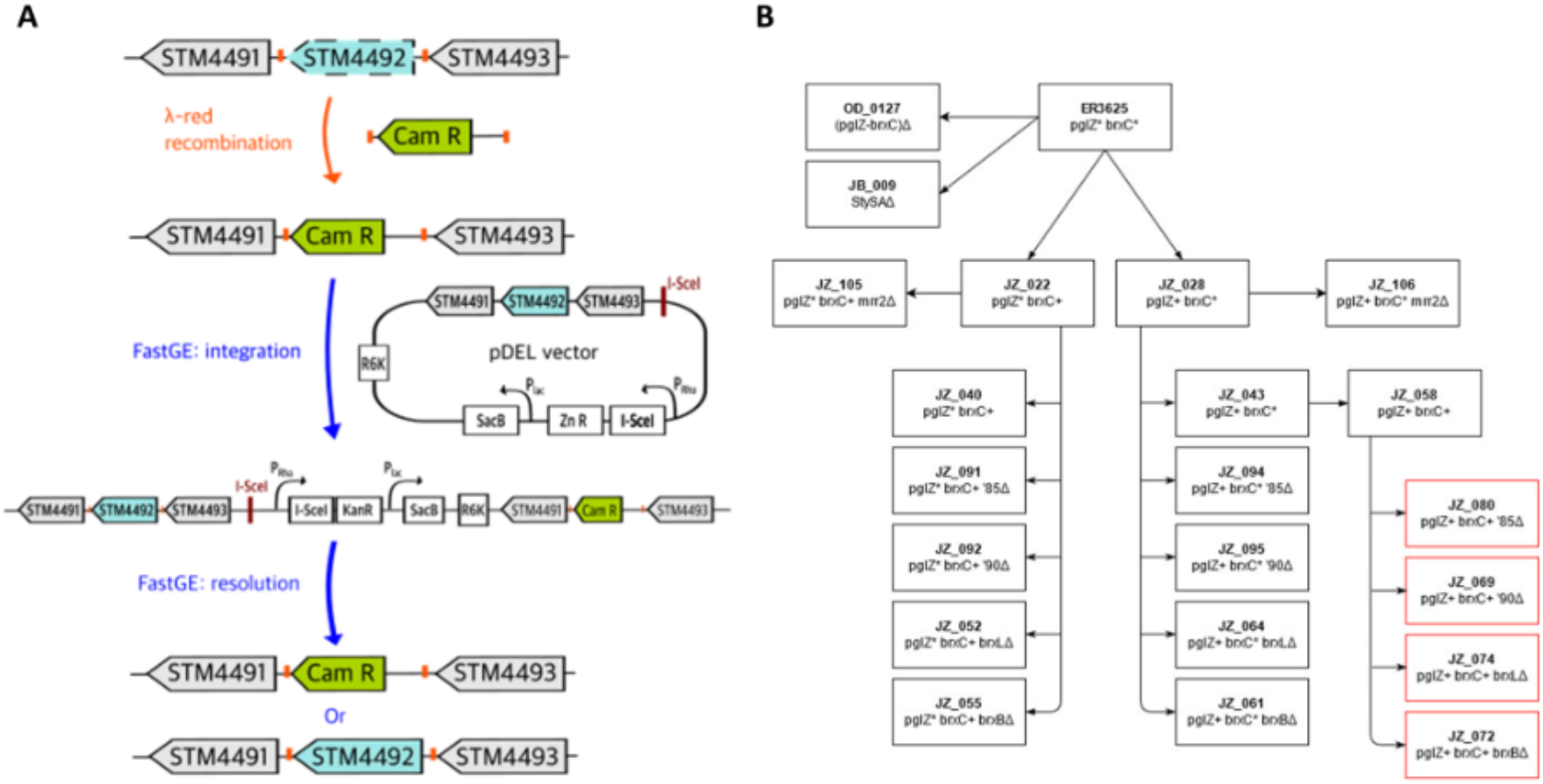
Engineering method and engineered strains. Panel A, Scarless engineering pipeline for replacement of mutated gene by wild type. Panel B, partial strain pedigree. Intermediate steps of panel A are omitted for clarity.

#### Cassette replacement of a gene deletion or gene segment

A *cat* cassette flanked by 36-50 bp of homology to the target gene was PCR-amplified from pKD3 and used to electroporate an intermediate host carrying pKD46, a *λ* Red-expressing thermosensitive vector [62]. Colony PCR with flanking gene-specific primers validated the construction (S3 File).

#### Replacement of the *cat* cassette with wild type sequence

The FAST-GE method was developed to rapidly engineer the genomes of both *E. coli* B and K12 strains with high efficiency by homologous recombination, with no residual extra sequence (“scar”) at the site of engineering. A single allele-exchange vector pDEL, was designed as compact as possible to maximize flexibility in application yield and fidelity. The pDEL vector contains several components involved in the promotion of recombination processes such as I-SceI endonuclease with its corresponding recognition site, and counterselection marker *sacB*. The unique double-strand break caused by I-SceI was shown to improve local recombination efficiency. Expression of *sacB* in the presence of sucrose is toxic to a variety of Gram-negative bacteria. To improve the overall efficiency of the protocol, *sacB* and gene coding for I-SceI endonuclease were placed under a *lac* and *rhaBAD* promoter respectively.

The original pDEL vector was constructed with a kanamycin resistance cassette as the primary selection marker. Since our host carries a kanamycin resistance cassette at the *mrr* locus, we designed two pDEL vectors with alternative antibiotic selection markers: zeocin and gentamycin (see sequences in S4 File). Both drug resistance cassettes were provided by Dr. Weigele. Zeocin is a bleomycin-like compound that kills both eukaryotic and prokaryotic cells by introducing lethal double-strand breaks in chromosomal DNA [64]. NEBuilder Hifi technology was used for construction of the pDEL-Zn (pOD003) and pDEL-Gm (pOD004) vectors (S2 File). Briefly, zeocin and gentamycin resistance cassettes were amplified by PCR with primers (oOD_070 and oOD_071) containing extensions homologous to the linearized pDEL vector without its kanamycin cassette (previously amplified with oOD_068 and oOD_069, S3 File), then assembled with the vector, transformed into *E. coli pir+* cells, selecting for the new drug resistance. Colony PCR with the set of primer oDO_072 and oDO_073 (S3 File) yields a product of 988 bp or 1.1 kb for pOD003 or pOD004 vector respectively. An amplicon of 48 bp is expected for the original pDEL vector (S4 File and S3 File).

This combination of methods was used successfully to engineer the strains characterized in this study for methylation and restriction activity (Figure 1B and S2 File). Cappable-seq data (Figure 2) were used in the strain design to limit as much as possible the disruption of TSS and transcription terminator structures (S2 Fig.).

#### Linear method for locus replacement

This method consists of amplifying a cassette and flanking homology regions to replace a genome segment with the cassette. Here we tested several lengths of homology regions (3, 5, 9 and 12 kb). 3 kb worked. First, we amplified the Zn cassette from pJBJZ_006 with oJB_018 and oJB_017 and the genomic flanking regions with dedicated primers (oJB_009/oJB_006 and oJB_007/oJB_010). These fragments were purified with NEB T1030 kit and ligated together using the NEBuilder^®^ Hifi DNA assembly with a ratio 1:1:1 (HR1: cassette: HR2) with a 30 min incubation. The assembly was then amplified using Q5^®^ High-Fidelity DNA Polymerase with an internal pair of primers (oJB_005/oJB_008) and repurified with NEB T1030 kit. Transformation was performed with 220 ng of the amplicons with Zn selection. Clones were verified by colony PCR.

### Phenotype tests

#### Growth media

Bacteria were grown in RB (10g soy peptone, 5g yeast extract, 5g NaCl per liter) or RBStrepKan (RB with Streptomycin (100 μg/mL) and Kanamycin (40 μg/mL) unless otherwise indicated.

#### Phage restriction tests

Bacteriophage L, with 13 sites for StySA [61] was used to test for StySA restriction, following the practice of the Colson and Ryu laboratories [40, 65]. Bacterial strains were grown in RB with antibiotic overnight at 37°C with agitation. The cultures were subcultured in new RB media without antibiotic and grown until exponential phase at 37°C.

Two-layer agar plates were used for the spot titer restriction test (S3 Fig.). The bottom layer is 1.5% agar (per liter: 15g of Bacto Agar BD Biosciences #214030, 10g of Bacto Tryptone BD Biosciences #211699, 5g of Bacto Yeast extract BD Biosciences #212720 and 5g of NaCl) and the top layer is an agar 0.7% (same recipe as the bottom layer but only 7 g of Bacto agar). Bacterial cultures were mixed with the top agar layer (56C) and poured on the bottom layer. The bacteriophage stocks (PH_JZ003 and PH_JZ006, see details in S2 File) were diluted from 10^-1^ to 10^-8^; 5 μl of each dilution was spotted on the plates; incubated at room temperature until dry, and incubated 18h at 37°C. Strains ER3625 and ER3649 were negative and positive controls. Plaques were counted on spots where they were well isolated.

#### Growth rate analysis

Growth rates were measured in 96 well plates with a plate reader (REF) with two technical replicates of each of three biological replicates. A single colony was inoculated in 1 ml RBStrepKan in a deep well plate, then incubated overnight at 37°C with shaking 200 rpm. A 96 well plate (Greiner Bio-One ref: 655892) was prepared with 200μl RBStrepKan per well and inoculated with 2μl of the overnight culture. The growth was monitored every 15 minutes for 15 hours, between each measure, the plate was shaken.

Growth rates were calculated from the raw data using a rolling regression from: https://padpadpadpad.github.io/post/calculating-microbial-growth-rates-from-od-using-rolling-regression/.https://padpadpadpad.github.io/post/calculating-microbial-growth-rates-from-od-using-rolling-regression/. A one-way ANOVA for normally distributed samples of non-equal variance was performed on the data to determine statistical significance of growth differences.

### Nucleic acid methods

#### Genomic DNA (gDNA) extraction, sequencing

Each strain was growth in RB with the appropriate antibiotics (see “Growth media” above) overnight at 37°C with 250 rpm agitation.

gDNA was extracted with the Monarch Genomic DNA purification kit (New England Biolabs; Ipswich, MA, USA) from 1ml of culture.

Libraries from these genomic DNAs were sequenced using the PacBio RSII or Sequel I sequencing platform. Briefly for RSII, SMRTbell libraries were constructed from genomic DNA samples sheared to between 10 and 20 kb using the G-tubes protocol (Covaris; Woburn, MA, USA), end repaired, and ligated to PacBio hairpin adapters. Incompletely formed SMRTbell templates and linear DNAs were digested with a combination of Exonuclease III and Exonuclease VII (New England Biolabs; Ipswich, MA, USA). The SMRTbell library was prepared according to PacBio sample preparation protocol sequenced with C4-P6 chemistry with a 300 min collection time.

For Sequel I libraries, SMRTbell libraries were constructed from genomic DNA samples following the PacBio protocol for Sequel using the kit 100-938-900. DNA qualification and quantification were performed using the Qubit fluorimeter (Invitrogen, Eugene, OR) and 2100 Bioanalyzer (Agilent Technology, Santa Clara, CA). The libraries were prepared for binding following the PacBio guidelines generated by SMRT Link and run on a Sequel I machine.

#### RNA extraction

For preculture three isolated colonies were grown in a 1ml RBStrepKan or RBStrepKan with Chloramphenicol (30 μg/mL) overnight at 37°C in a deep well with breathable cover tape. The cultures were subcultured the next day in 25ml RBStrepKan (no chloramphenicol) at 37°C with 250 rpm agitation. Cells were harvested when OD 600 nm reached ~0.3 by centrifugation 10 min at 4°C. Pellets were resuspended in 100μl of cold 0.1X PBS.

RNA was extracted using the Qiagen RNA extraction kit following the classical protocol for bacterial RNA. The eluted RNA was then treated with DNAse I (NEB) and then cleaned and concentrated using a classical phenol-chloroform RNA extraction. RNA was stored at −80°C.

#### TSS determination using Cappable-seq method

The TSS Cappable-seq libraries were prepared following the recommendation of the protocol from reference [47] starting from 2μg of RNA in duplicates and controls. The libraries were run with a MiSeq with 1 × 75bp insert size using V3 Illumina platform.

The analysis was run in command line from the raw data using the script available at https://github.com/Ettwiller/TSS/.

Quantitation of *brxX* transcription employed qPCR. Primers used are listed in S3 File. The Lunascript RT Supermix kit E3010 was used for random conversion of 500 ng of extracted RNA to cDNA per samples (three biological replicates per strain) following the kits guidelines. The no-RT reactions were run on the same plate with the same RNAs. qPCR was then run with 2 primer sets, *yceB* (oJZ_116 and oJZ_117) and *brxX*(oJZ_150 and oJZ_151) using the Luna Universal qPCR Master Mix Protocol (M3003) with 1μl of cDNA per well. For each primer pair a standard curve was run on the same plate as the sample with 1, 0.1, 0.01, 0.001 and 0.0001ng and water only. All combinations were run in duplicate on the same plate. The plates were sealed and centrifuged for 30s and then run 39 cycles on the same CFX96 Touch Bio-Rad machine following the Luna kit recommended cycle steps.

### Bioinformatic methods

#### Annotation of predicted functional domains and transcription signals

Predicted functional protein domains were annotated using Genbank-assigned protein IDs listed in S1 File for LT2 and for ER3625. The NCBI protein IDs are automatically annotated with “regions” that correspond to Conserved Domain Database [66] concise predictions. Additional automated domain annotations were generated and visualized by submission to InterPro using the Geneious implementation of InterProScan as described in S1 File. Manual search of the Conserved Domain Database with “Full results” instead of “Concise results” also elicits annotations compatible with InterPro.

Potential transcription start sites were documented experimentally as described below (CappableSeq). Rho-independent transcription terminators were predicted using TransTerm HP algorithm version 2.09 ([67, 68]), then curated manually.

#### Sequence verification and methylation analysis

The BREX engineered locus sequence was verified in two ways. At first, PCR were run with primers (oJZ_241/242, see S3 File) with LongAmp polymerase (New England Biolabs; Ipswich, MA, USA), purified and sequenced (Sanger sequencing?). The other method was to check the sequences of the modified locus with the PacBio sequencing reads from the methylation analysis.

DNA motifs and degree of modification were generated using InterPulse Duration (IPD) Ratios analyzed with RS_Modification_and_Motif_Analysis from PacBio as in [69, 70]. PBMotStat, a component of REBASE TOOLS [71] was used to calculate the % of methylated sites with IPD > 2 for specific sites.

For the partially methylated strains, a .gff file of the methylation level of each single base of the genome was downloaded from SMRT Link, filtered to keep only significantly methylated sites, and uploaded in Geneious Prime on the reference genome (File S6).

#### Global and local transcriptome analysis by RNAseq

RNAseq libraries were prepared with the “protocol for library preparation of Intact RNA using NEBNext rRNA depletion kit (Bacteria) (NEB#E7850, NEB#78860) and NEBNext Ultra II directional RNA library prep kit for Illumina (NEB#E7760, NEB#E7765)” from 250ng of sample. Libraries were barcoded with dual index from NEB Multiplex Oligos for Illumina (96 Unique Dual Index Primer Pairs set 4, E6446S). The sequencing of the pooled libraries was performed on a NextSeq apparatus

The raw reads were downloaded on Galaxy (Version 0.4.3.1) to perform the QC, mapping, and feature assignment (S4 Fig.). After a first QC step to assess the quality of the reads, they were trimmed using TrimGalore (Version 0.4.3) with parameters: -q 20 -s 3 -e 0.1 –length 20 and then a second QC steps was performed to verify the read quality after trimming. The trimmed reads were mapped with bowtie2 (with default parameters except –fr for upstream/downstream mate orientations) to the ER3625 genome (NCBI CP067091-CP067092 and [61, 72]. The alignment was then used to count the number of fragments mapped to CDS and rRNA features using in parallel Featurecounts [72] (parameters: -s stranded (reverse)) and htseq (Version 0.9.1) with parameters: --mode Union –stranded reverse –minaqual 10 --idattr locus_tag –nonunique no.

The downstream part of this analysis was performed using the Deseq2 R package [73] in command line. The parameters were set as: lfcShrink = apeglm, padj < 0.05 and log2FoldChange > |1.5|. UpsetR was used for visualization (Figure 6) [74]. Intersect (ref) was used to generate lists of features common to chosen strain sets used in S5 File.

## Acknowledgements

Eddie Gilcrease, Sherwood Casjens and Peter Weigele have been a wonderful source of knowledge about the bacteriophage L and its use in *Salmonella* strains.

Bo Yan, Brad Langhorst, Zhiyi Sun and Amogh Changavi for bioinformatic advice and training.

Julie Beaulieu and Jim Samuelson the pDEL vector and protocol and submission to Addgene.

Barry Stoddard, Rick Morgan and Tim Blower for sharing ideas and discussions BREX systems and their contexts.

New England Biolabs Research: Protein Expression and Modification Division and E. coli club for discussions and “annoying” questions that helped build the project.

Donald Comb in memoriam for fostering basic research because it’s exciting.

## Supplementary Materials

S2_Fig_Sup_data_KO_segments.docx. The figure shows segments missing in *cat*::KO constructions related to TSS and TTS; the document includes a legend

S3_Fig_colonies_restriction.docx. The figures show A. colony morphology for all strains; B. Illustrative phage titer figure and table of restriction magnitude for all strains. The document includes a legend.

S4_Fig_Galaxy_pipeline.docx. The document includes a legend with link to Galaxy public web site.

S1_File_Fig_Table_protocol_Domain_predictions_six_StySA_components.docx. The document includes a figure illustrating the domain hits identified, a table of locus_tags and protein_IDs used for search, and a protocol used to generate the hit list and visualization figure.

S2_File_plasmid_strain_phage.xlsx. Strain list with genotype and construction steps; plasmids with description and availability; phage with source and identity and RM phenotype of last host.

S3_File_Sup_Data_primers.xlsx. Oligonucleotide sequences used for plasmid construction, strain engineering, validation of sequences and qPCR.

S4_File_ Sup_Data_Sequences_cassettes_plasmids.docx. This contains sequences highlighted in color to navigate the locations of primers, coding sequences, promoters, terminators, replicons, and protein binding sites.

S5_File_RNAseq_hits_annotated_and_coded.

S6_File_partial_methylation.zip. This file combines a reference sequence and gff file for modification of independently constructed isogenic strains JZ_055, JZ_056, JZ_057.

S7_File_Potential_BrxL_activities.docx. This includes three figures and extended discussion of activities of other proteins with domain hits like that of the BrxL C-terminal domain.

S8_File_Sup_Data_SRA.xlsx. This describes the contents of SRA files: strain, NEB strain ID if applicable, Biosample Name (NCBI-assigned), analysis type (RSII, Sequel, Cappable-seq, NextSeq RNAseq)

S9_File_Transcript_Termination_Sites.docx. This document contains four figures: strategy, and three IGV sequence read alignment screenshots at different magnifications. No legend at present.

S10_File_ Protocol_for_adding_annotations_to_RNASeq_output_files

## Data availability (this is in a box at one side)

All the data can be found in the BioProject PRJNA605961. All the raw data are available under the SRA (See S8 File Sup_Data_SRA,xlsx).

All the Salmonella strains and phages are available from New England Biolabs. The plasmids are available at Addgene.

